# Nintedanib ameliorates experimental pulmonary arterial hypertension via inhibition of endothelial mesenchymal transition and smooth muscle cell proliferation

**DOI:** 10.1101/584110

**Authors:** Takeo Tsutsumi, Tetsutaro Nagaoka, Takashi Yoshida, Lei Wang, Sachiko Kuriyama, Yoshifumi Suzuki, Yuichi Nagata, Norihiro Harada, Yuzo Kodama, Fumiyuki Takahashi, Yoshiteru Morio, Kazuhisa Takahashi

## Abstract

Neointimal lesion and medial wall thickness of pulmonary arteries (PAs) are common pathological findings in pulmonary arterial hypertension (PAH). Platelet-derived growth factor (PDGF) and fibroblast growth factor (FGF) signaling contribute to intimal and medial vascular remodeling in PAH. Nintedanib is a tyrosine kinase inhibitor whose targets include PDGF and FGF receptors. Although the beneficial effects of nintedanib were demonstrated for human idiopathic pulmonary fibrosis, its efficacy for PAH is still unclear. Thus, we hypothesized that nintedanib is a novel treatment for PAH to inhibit the progression of vascular remodeling in PAs. The inhibitory effects of nintedanib were evaluated both in endothelial mesenchymal transition (EndMT)-induced human pulmonary microvascular endothelial cells (HPMVECs) and human pulmonary arterial smooth muscle cells (HPASMCs) stimulated by growth factors. We also tested the effect of chronic nintedanib administration on a PAH rat model induced by Sugen5416 (a VEGF receptor inhibitor) combined with chronic hypoxia. Nintedanib was administered from weeks 3 to 5 after Sugen5416 injection, and pulmonary hemodynamics and PAs pathology were evaluated. Nintedanib attenuated the expression of mesenchymal markers in EndMT-induced HPMVECs and HPASMCs proliferation. Phosphorylation of PDGF and FGF receptors was augmented both in both intimal and medial lesions of PAs. Nintedanib blocked these phosphorylation, improved hemodynamics and reduced vascular remodeling involving neointimal lesions and medial wall thickening in PAs. Additionally, expressions Twist1, transcription factors associated with EndMT, in lung tissue was significantly reduced by nintedanib. These results suggest that nintedanib may be a novel treatment for PAH with anti-vascular remodeling effects.

## Introduction

The pathogenesis of pulmonary arterial hypertension (PAH) involves abnormal vasoconstriction and vascular remodeling in pulmonary arteries (PAs). Various stimuli, including hemodynamic stress, hypoxia, and inflammation, alter endothelial cell functions and cause an imbalance in endothelial cell-derived vasoconstrictive and vasodilative factors (1). Endothelin-1, prostacyclin, and nitric oxide are major regulators of vascular tone, and several types of vasodilators for PAH have been developed that target these signaling pathways. Although the prognosis of PAH has been remarkably improved by these vasodilators (2, 3), there is still no drug that targets vascular remodeling in PAH.

Pulmonary arterial remodeling plays an essential role in the progression of PAH, especially in the late phase of disease. Previous studies have suggested that platelet derived growth factor (PDGF) and fibroblast growth factor (FGF) signaling contribute to vascular cell proliferation, migration, differentiation, and apoptosis (4). Expressions of PDGF-A, PDGF-B, PDGF receptor-α, and PDGF receptor-β are increased in the PAs of PAH patients compared with those of healthy patients (5). Previous reports have also demonstrated elevated basic FGF concentration in the plasma from PAH patients (6). FGF2 signaling mediated by FGF receptor 1 (FGFR1) promotes proliferation of vascular endothelial cells (7) and smooth muscle cells (8) in human PAH. Additionally, endothelial mesenchymal transition (EndMT) has been suggested to contribute to the progression of occlusive neointimal lesions in PAs, which is a characteristic vascular abnormality in PAH (9, 10). Vascular endothelial cells acquire a mesenchymal phenotype following exposure to several cytokines and growth factors, including transforming growth factor β (TGF-β), PDGF, and FGF which are induced by shear stress, hypoxia, and inflammation (10). EndMT-modified endothelial cells acquire additional characteristics of mesenchymal cells, and recent reports have indicated a critical role of EndMT in the development of PAH-specific neointimal lesions in PAs (11–13). Although both medial wall thickness and neointimal lesions in PAs are common pathological findings in PAH, a greater contribution of neointimal lesions in the elevation of pulmonary arterial pressure was reported in experimental PAH (14), suggesting an important involvement of EndMT.

Nintedanib is a triple tyrosine kinase inhibitor (TKI) of PDGF, FGF, and vascular endothelial growth factor (VEGF) receptors (15). A recent phase III clinical study showed that nintedanib prevented both the progression of restrictive pulmonary impairment and the acute exacerbation of idiopathic pulmonary fibrosis (IPF), with good tolerability (16). Thus, nintedanib was approved globally for the treatment of IPF. However, the efficacy of nintedanib to treat PAH by inhibiting PDGF and FGF signaling is still uncertain. Based on this background information, we hypothesized that nintedanib is a novel and beneficial treatment for PAH by targeting vascular remodeling including neointimal lesion and medial wall thickening in PAs.

## Materials and Methods

Primary human pulmonary microvascular endothelial cells (HPMVECs; product code. CC-2527, lot no. 547317, 560121, and 621712) and primary human pulmonary arterial smooth muscle cells (HPASMCs; product code. CC-2581, lot no. 029837, 407340, and 559495) were obtained from Lonza (Basel, Switzerland). All experimental and surgical procedures were approved by the Institutional Committee for the Use and Care of Laboratory Animals in Juntendo University (Tokyo, Japan), in accordance with the U.S. National Institutes of Health Guide for the Care and Use of Laboratory Animals.

### EndMT of HPMVECs

HPMVECs were cultured in Microvascular Endothelial Cell Growth Medium-2 (EGM™-2MV, Lonza). TGF-β2 (2.5 ng/mL), tumor necrosis factor (TNF)-α (2 ng/mL), and interleukin (IL)-1β (4 ng/mL) were used for the induction of EndMT as previously described (17). The cytokines were added to the medium with or without a 3-h preincubation with nintedanib (1 μM). This concentration of nintedanib had been confirmed to have maximum inhibitory effects in preliminary experiments. The expressions of von Willebrand factor (vWF) and CD31 proteins, which are endothelial markers, as well as fibronectin and collagen 1 proteins, which are mesenchymal markers, were analyzed at 48 h after stimulation by western blotting to confirm EndMT of the HPMVECs.

### Proliferation assay of HPASMCs

HPASMCs were cultured in Smooth Muscle Growth Medium-2 (SmGM™-2, Lonza). The proliferation of HPASMCs was evaluated using both the Cell Counting Kit-8 (CCK-8) and Bromodeoxyuridine (BrdU) ELISA kit as previously described, respectively (18, 19). HPASMCs were treated with multiple growth factors, specifically 5% fetal calf serum, PDGF-BB (30 ng/mL), FGF2 (2 ng/mL), epidermal growth factor (EGF) (0.5 ng/mL), and insulin-like growth factor-1 (IGF-1) (0.5 μg/mL) with or without imatinib (3 μM) or nintedanib (0.3 μM). The viability of HPASMCs was evaluated time-dependently after stimulation. In advance of this assay, we examined the concentration-dependent inhibitory effects of nintedanib and imatinib, the latter of which is another TKI for PDGF signaling, on the proliferation of HPASMCs induced by PDGF-BB. Based on these preliminary results, we selected 0.3 μM and 3 μM of nintedanib and imatinib, respectively, as concentrations that produced maximum inhibitory effects. LDH assay was also performed to assess the cell toxicity of nintedanib and imatinib as previously described (20). Value of LDH in the medium with nintedanib and imatinib were normalized by that in the control medium without TKI. The protein expression of extracellular signal-regulated kinase (ERK)1/2 and AKT with or without phosphorylation, which are downstream effectors of PDGF and FGF signaling, was also evaluated by western blotting at 2 h after stimulation of the HPASMCs.

### Preparation of pulmonary hypertensive rats

The PAH rat model was established using Sugen 5416 (VEGF receptor-1,-2 inhibitor) and chronic hypoxic exposure as previously described (14, 21). Briefly, adult male Sprague Dawley rats (150 – 180 g) were injected with Sugen 5416 (20 mg/kg; Cayman Chemical Co, MI) subcutaneously and exposed to hypobaric hypoxia (360 mmHg, 10% O_2_) for 3 weeks. After returning to normoxic conditions (760 mmHg, 21% O_2_), the rats were analyzed immediately [SuHx(3W) group] or were chronically treated with either vehicle (0.5% hydroxyethylcellulose) [SuHx(5W) group] or nintedanib [50 mg/kg/day, SuHx(5W) + Nin group] by oral gavage for 2 weeks. The dose of nintedanib was determined based on previously published experiments (15). Control rats received a single vehicle injection and were exposed to normoxic conditions for 5 weeks, with vehicle or nintedanib (50 mg/kg/day) gavage from 3 to 5 weeks [Nx(5W)] and [Nx(5W) + Nin] groups, respectively. Hemodynamic measurements were performed at 3 or 5 weeks after vehicle or Sugen 5416 injection.

### Pulmonary hemodynamic measurements

Pulmonary hemodynamic measurements by right heart catheterization were performed as previously described (22). Briefly, all rats were anesthetized with pentobarbital sodium (30 mg/kg intraperitoneal). A polyvinyl catheter was inserted into the right ventricle (RV) via the right jugular vein for the measurement of RV systolic pressure (RVSP) with the PowerLab data acquisition system (AD Instruments, CO). Systemic systolic arterial pressure (SAP) and heart rate (HR) were continuously monitored, and rats with an HR of consistently less than 300 beats/min were excluded. After the hemodynamic measurements, all rats were euthanized by an overdose of pentobarbital sodium, and their hearts and lungs were collected for RV / left ventricle (LV) + septum weight ratio (RV/LV+S) measurements to show RV hypertrophy and for histological evaluations. The right lungs were stored for protein measurement, and the left lungs were inflated with 10% buffered formalin at a constant pressure of 20 cm H_2_O for histological analyses. Fixation was allowed to proceed overnight.

### Morphological analysis of PAs

The inflated and fixed left lungs were embedded in paraffin. All sections were cut at 5-μm thickness and were stained with elastic Van Gieson stain. A quantitative analysis of PA luminal obstruction was performed as described previously with minor modifications (14). We counted between 100 and 200 small PAs [outer diameter (OD) < 200 μm] per whole left lobe at ×400 magnification using an image analysis system (KS400; Carl Zeiss Imaging Solutions, Germany) in a blind manner. The medial-wall thickness was measured for the arteries with an OD between 50 and 200 μm. The distance between the internal and external elastic laminae was expressed as the medial thickness/OD. Vessels with an OD < 50 μm were used for the assessment of occlusive neointimal lesions and were scored as follows: no evidence of occlusive neointimal formation (grade 0), partial luminal occlusion (≤ 50%; grade 1), and severe luminal occlusion (> 50%; grade 2).

### Immunohistochemistry of PAs

After deparaffinization with xylene, rehydration, and antigen retrieval by heating in citrate buffer (pH 6), immunohistochemistry was performed with an anti-FGFR1 (phospho Y654) antibody (ab59194, Abcam, Cambridge, UK), anti-PDGF receptor-β (phospho Y1020) antibody (ab16868, Abcam, Cambridge, UK). The signals were detected using VECTASTAIN^®^ elite ABC rabbit, mouse and goat IgG kits (#PK-6101, 6102, 6105, Vector Laboratories, CA). In each whole left lobe, 100 arteries across serial sections were examined to evaluate the expression of each receptor.

### Expression of Twist1 in rat lung tissues

The expression of Twist1 protein, which is an EndMT-related transcription factor (11), was evaluated by western blotting in lung tissues from Nx(5W), SuHx(5W), and SuHx(5W) + Nin groups to assess the contribution of EndMT in the vascular remodeling of PAs in rat PAH and to assess the anti-EndMT effect of nintedanib.

### Western blotting analysis

HPASMCs, HPMVECs, and lung tissues were lysed in radioimmunoprecipitation assay (RIPA) buffer containing protease and phosphatase inhibitors, and the lysates were subjected to western blotting, as described previously (23). Transferred membranes were allowed to react with anti-vWF polyclonal antibody (1:1,000; sc14014, Santa Cruz Biotechnology), anti-CD31 monoclonal antibody (1:2,000; bba7, R&D Systems, MN), anti-fibronectin monoclonal antibody (1:2,000; sc59826, Santa Cruz Biotechnology), anti-Collagen 1 polyclonal antibody (1:1,000; ab34710, Abcam), anti-p44/42 MAPK (Erk1/2) Antibody (1:2,000; #9102, Cell Signaling Technology), anti-Phospho-p44/42 MAPK (Erk1/2) (Thr202/Tyr204) Antibody (1:2,000; #9101, Cell Signaling Technology), anti-AKT polyclonal antibody (1:2,000; #9272, Cell Signaling Technology), anti-phospho-AKT (Ser473) polyclonal antibody (1:2,000; #9271, Cell Signaling Technology), anti-Twist1 polyclonal antibody (1:1000; ab50581, Abcam), or anti-β-actin monoclonal antibody (1:10,000; A5441, Sigma, MO). Western blot signals were acquired using a Fuji ImageQuant™ LAS-4000 fluorescence imager and quantified using the Multi Gauge image analysis software (Fujifilm Corporation, Tokyo, Japan). The densitometric signal of each protein was normalized to that of β-actin.

### Statistical analysis

Data are presented as means ± SE. Statistical analysis was performed using one-way ANOVA (Prism 6; GraphPad Software, La Jolla, CA, USA). Differences were considered significant at *P* < 0.05.

## Results

### Inhibitory effect of nintedanib on EndMT of HPMVECs

We confirmed decreased expression of vWF and CD31 proteins and increased expression of fibronectin and collagen 1 proteins by western blotting. Nintedanib attenuated the upregulation of mesenchymal markers in the stimulated HPMVECs, but did not prevent the downregulation of endothelial markers (Figure 1A, B). After stimulation with TGF-β2, TNF-α, and IL-1β, the morphology of HPMVECs changed from a cobblestone to spindle-shaped morphology, and nintedanib tended to prevent its change. (Figure 1C).

**Figure 1.**
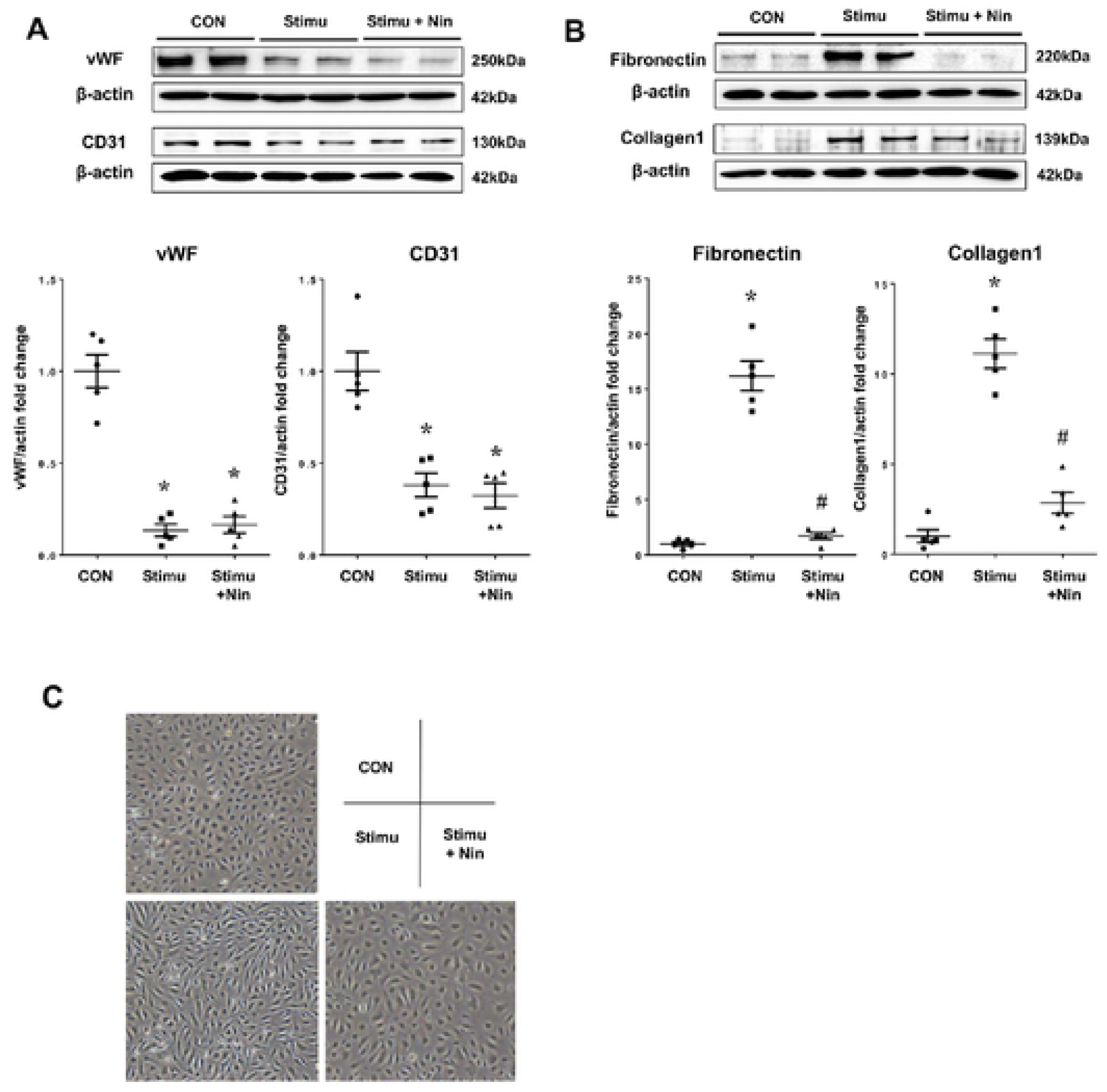
Inhibitory effect of nintedanib on endothelial mesenchymal transition of human pulmonary microvascular endothelial cells. Inhibitory effect of nintedanib (Nin) on endothelial–mesenchymal transition (EndMT) of human pulmonary microvascular endothelial cells (HPMVEC). EndMT was induced by stimulation with TGF-β2, TNF-α, and IL-1β (Stimu). Unstimulated cells are denoted as CON. Expression of (A) von Willebrand factor (vWF) and CD31 proteins, which are endothelial markers, and (B) fibronectin and collagen 1 proteins, which are mesenchymal markers. * p<0.01 vs. CON. # p<0.001 vs. Stimu. (C) Representative morphology of HPMVECs in Con, Stimu, and Stimu+Nin condition.

### Inhibitory effect of nintedanib on proliferation of HPASMCs

The proliferation of HPASMCs induced by multiple growth factors (PDGF-BB, FGF2, EGF, and IGF) was significantly greater than that without stimulation in CCK-8 and BrdU assays, and nintedanib significantly inhibited this proliferation at 24 and 48 h after stimulation. Moreover, the inhibitory effect of nintedanib was significantly greater than that of imatinib at 48 h after stimulation in CCK-8 assay (Figure 2A, 2B). Values of LDH in the medium after treatment with imatinib and nintedanib were similar between all groups (Figure 2C). The phosphorylation of ERK1/2 and AKT was increased in the HPASMCs stimulated with multiple growth factors, and nintedanib remarkably prevented these phosphorylations. This preventive effect of nintedanib was significantly greater than that of imatinib on the phosphorylated AKT (Figure 2D).

**Figure 2.**
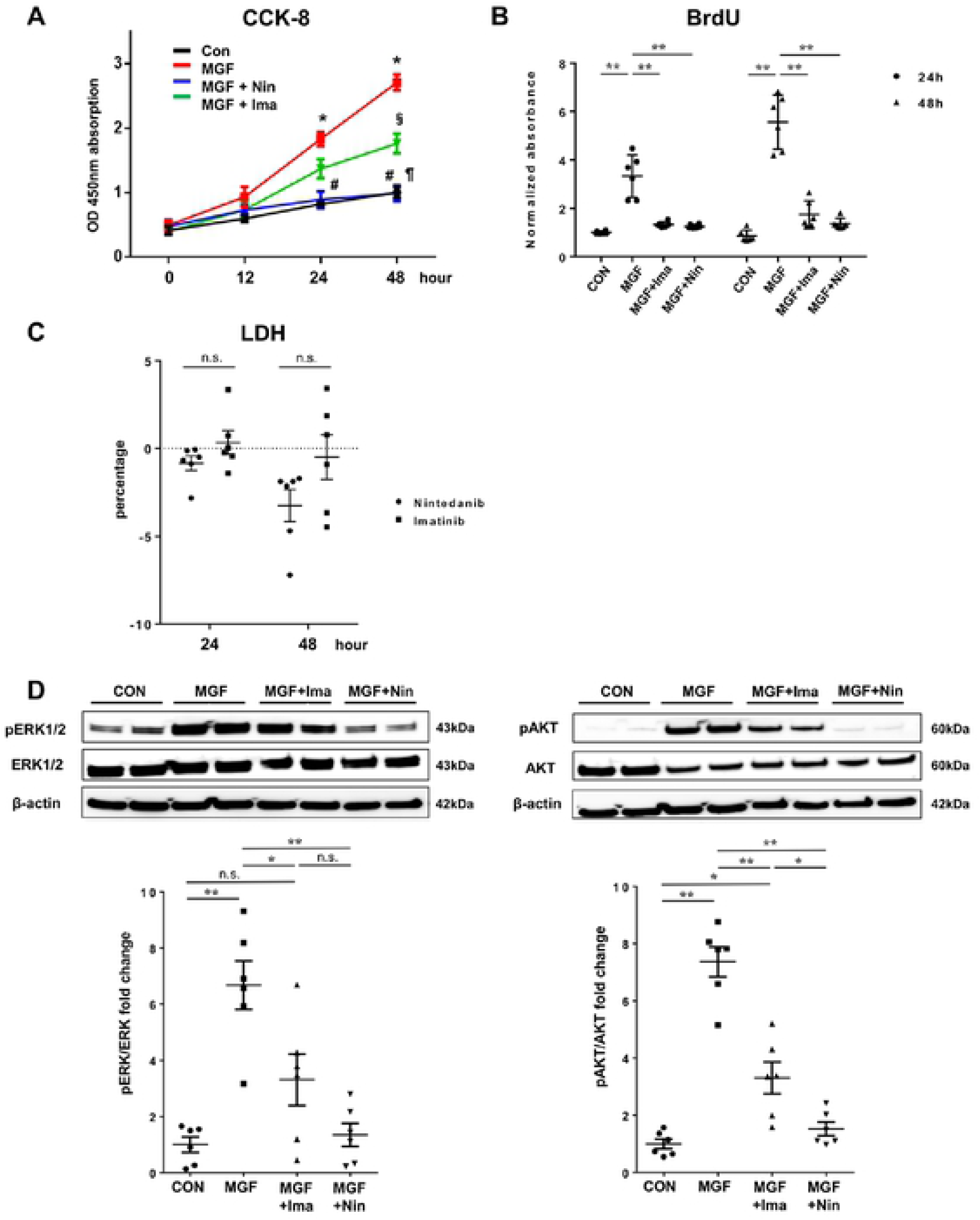
Inhibitory effect of nintedanib on proliferation of human pulmonary arterial smooth muscle cells. (A) Inhibitory effect of nintedanib (Nin) and imatinib (Ima) on proliferation of human pulmonary arterial smooth muscle cells (HPASMCs) using the CCK-8 assay. Cell viability of HPASMC was assessed at 0, 12, 24, and 48 h after stimulation with multiple growth factors (MGF, platelet derived growth factor-BB + fibroblast growth factor 2 + epidermal growth factor + insulin-like growth factor-1). Con: control HPASMC without MGF stimulation. Values are means ± SE (n=5). * p<0.001 MGF vs. Con. § p<0.01 MGF vs. MGF + Ima. # p<0.01 MGF vs. MGF + Nin. ¶ p<0.05 MGF + Nin vs. MGF + Ima. (B) Inhibitory effect of Nin and Ima on proliferation of HPASMC using the BrdU assay. Cell viability of HPASMC was assessed at 24 and 48 h after stimulation with MGF. ** p<0.001. Plotted values are means ± SE (n=6). (C) Cell toxicity of Nin and Ima on HPASMC using the LDH assay. Values of LDH were assessed at 24 and 48 h after treatment of Nin and Ima. Value of LDH in the medium with Nin and Ima were normalized by that in the control medium. Plotted values are means ± SE (n=6). (D) Expressions of ERK1/2, phosphorylated ERK1/2 (pERK1/2), AKT, and phosphorylated AKT (pAKT) protein with or without Nin or Ima by western blotting. Representative blots are shown in the upper panels. Densitometric signals of the phosphorylated protein were normalized to the corresponding non-phosphorylated protein. Plotted values are means ± SE (n=6). * p<0.01. ** p<0.001. n.s. not significantly.

### Chronic treatment with nintedanib in PAH rats

The rats were divided into 5 groups as follows: 5-week normoxia + vehicle group [Nx(5W)], 5-week normoxia + nintedanib group [Nx(5W) + Nin)], Sugen 5416 + hypoxia + vehicle group [SuHx(3W) and SuHx(5W)], and Sugen 5416 + hypoxia + nintedanib group [SuHx(5W) + Nin] (Figure 3A). Three weeks after Sugen 5416 injection and hypoxic exposure, the RVSP and RV/LV+S were already higher in the SuHx(3W) group (81.3±2.7 mmHg and 0.688±0.034, respectively) than in the Nx(5W) group (22.4±1.1 mmHg and 0.296±0.014, respectively). After 2 weeks of treatment with nintedanib [SuHx(5W) + Nin], the RVSP and RV/LV+S were significantly reduced (50.4±7.2 mmHg and 0.546±0.029, respectively) compared with those of the SuHx(5W) group (81.8±6.6 mmHg and 0.728±0.023, respectively). Treatment with nintedanib under normoxic condition [Nx(5W) + Nin)] caused no effect on pulmonary hemodynamics (Figure 3B). Chronic treatment with nintedanib also did not affect the systemic SAP, HR, or body weight (Figure 3C). The values of RVSP and RV/LV+S in the SuHx(5W) rats are consistent with those of a previous study (14).

**Figure 3.**
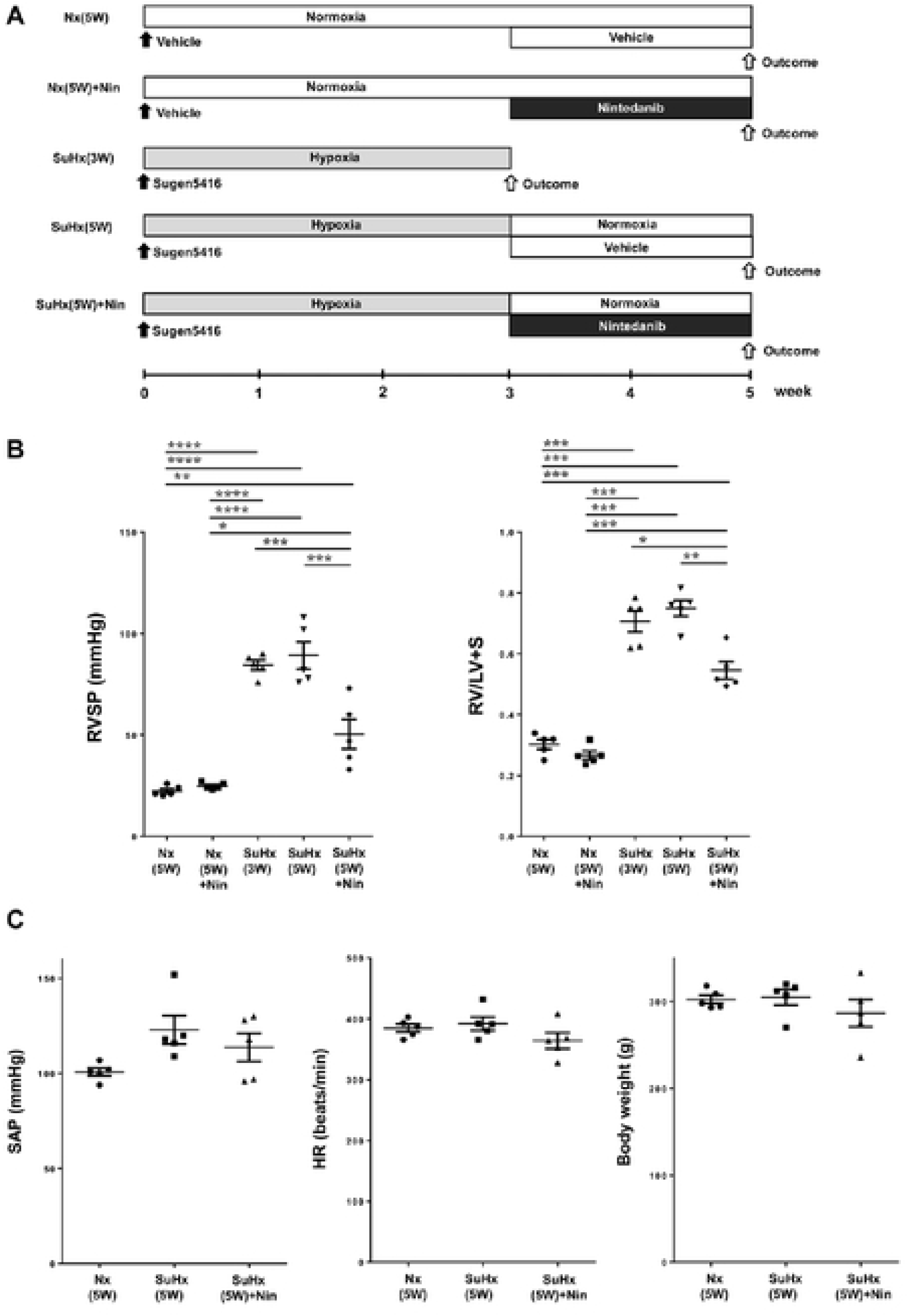
Effect of chronic nintedanib for the development of pulmonary arterial hypertension in rat. (A) *In vivo* experimental timeline. Outcomes include measurement of pulmonary hemodynamics and right ventricle hypertrophy, and/or morphometric analysis of pulmonary arteries. Nx: control rat with normoxic condition. SuHx: single injection of Sugen 5416 with chronic hypoxic exposure. W: week. (B) Chronic effect of nintedanib in Su/Hx rat. Examinations of right ventricle systolic pressure (RVSP) and right ventricle weight / (left ventricle and septal weight) ratio (RV/LV+S) in normoxic rats at 5 weeks [Nx(5W)], Nx rat with nintedanib treatment from weeks 3 to 5 [Nx(5W) + Nin], Su/Hx rat at 3 weeks [SuHx(3W)] or at 5 weeks [SuHx(5W)] after Sugen 5416 injection, and SuHx rats with nintedanib treatment from weeks 3 to 5 [SuHx(5W) + Nin]. (C) Examinations of systemic systolic arterial pressure (SAP), heart rate (HR), and body weight in Nx(5W), SuHx(5W), and SuHx(5W) + Nin rats. Plotted values are means ± SE (n=5). * p<0.05. ** p<0.001. *** p<0.0005, **** p<0.0001.

The medial wall thickness of PAs (50 to 200 μm OD) in the [SuHx(5W)+Nin] group was significantly reduced compared to that in the SuHx(3W) and SuHx(5W) groups. The grade 1 and grade 2 neointimal occlusive lesions in the small PAs (< 50 μm OD) were also significantly lower in [SuHx(5W) + Nin] than in SuHx(3W) and in SuHx(5W). Although the medial wall thickening and grade 1 neointimal occlusive lesions were reversed by treatment with nintedanib compared with the SuHx(3W) values, nintedanib treatment lessened, but did not reverse, the development of grade 2 neointimal occlusive lesions (Figure 4A, B). Both medial wall thickness and neointimal lesions with the severe PH group tended to be progressive compared to those of the less severe PH group.

**Figure 4.**
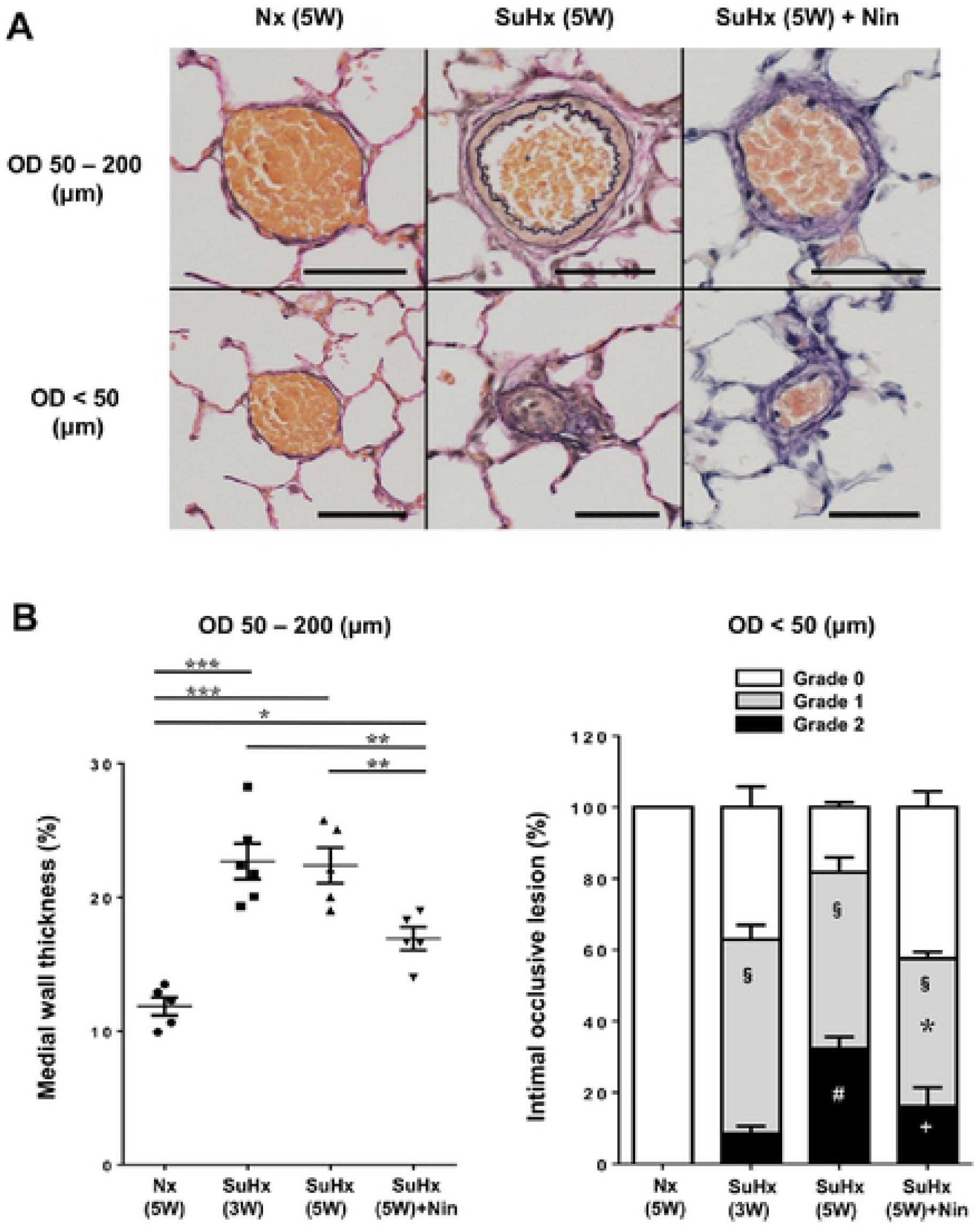
Effect of chronic nintedanib for the remodeling of small pulmonary arteries. (A) Representative elastic Van Gieson staining of pulmonary arteries in pulmonary normotensive control rats [Nx(5W)], Sugen 5416/hypoxic rats at 5 weeks after Sugen 5416 injection [SuHx(5W)], and SuHx rats with nintedanib treatment from weeks 3 to 5 [SuHx(5W) + Nin] (200×). Pulmonary arteries were divided in 50 to 200 μm outer diameter (OD) and < 50 μm OD for analysis. Scale bars indicate 50 μm (OD 50-200) and 10 μm (OD < 50). (B) Quantitative analysis of medial wall thickness in pulmonary arteries (left; 50 to 200 μm OD) and intimal occlusive lesions in smaller pulmonary arteries (right, < 50 μm OD) in pulmonary normotensive control rats at 5 weeks [Nx(5W)], Sugen 5416/hypoxic rats at 3 weeks [SuHx(3W)] or at 5 weeks [SuHx(5W)] after Sugen 5416 injection, and SuHx rats with nintedanib treatment from weeks 3 to 5 [SuHx(5W) + Nin]. Grade 0 (no lumen occlusion; white), grade 1 (<50% occlusion; gray), grade 2 (>50% occlusion; black). Plotted values are means ± SE (n=5-6). * p<0.05. ** p<0.001. *** p<0.0001 (left). **§** P<0.0001 vs. Nx(5W). * p<0.05 vs. SuHx(3W). # p<0.001 vs. SuHx(3W). + p<0.05 vs. SuHx(5W) (right).

### Phosphorylation of FGF and PDGF receptors in PAs

The expression of phosphorylated FGFR1 and PDGF receptor-β were significantly increased not only in PAs with medial wall thickening (OD 50–200 µm), but also in PAs with occlusive neointimal lesions (OD < 50 µm), in contrast to the low levels in the PAs of Nx(5W) lungs. The phosphorylation of FGFR1 and PDGF receptor-β in PAs was significantly decreased by 2-week treatment with nintedanib [SuHx(5W) + Nin] (Figure 5A, B).

**Figure 5.**
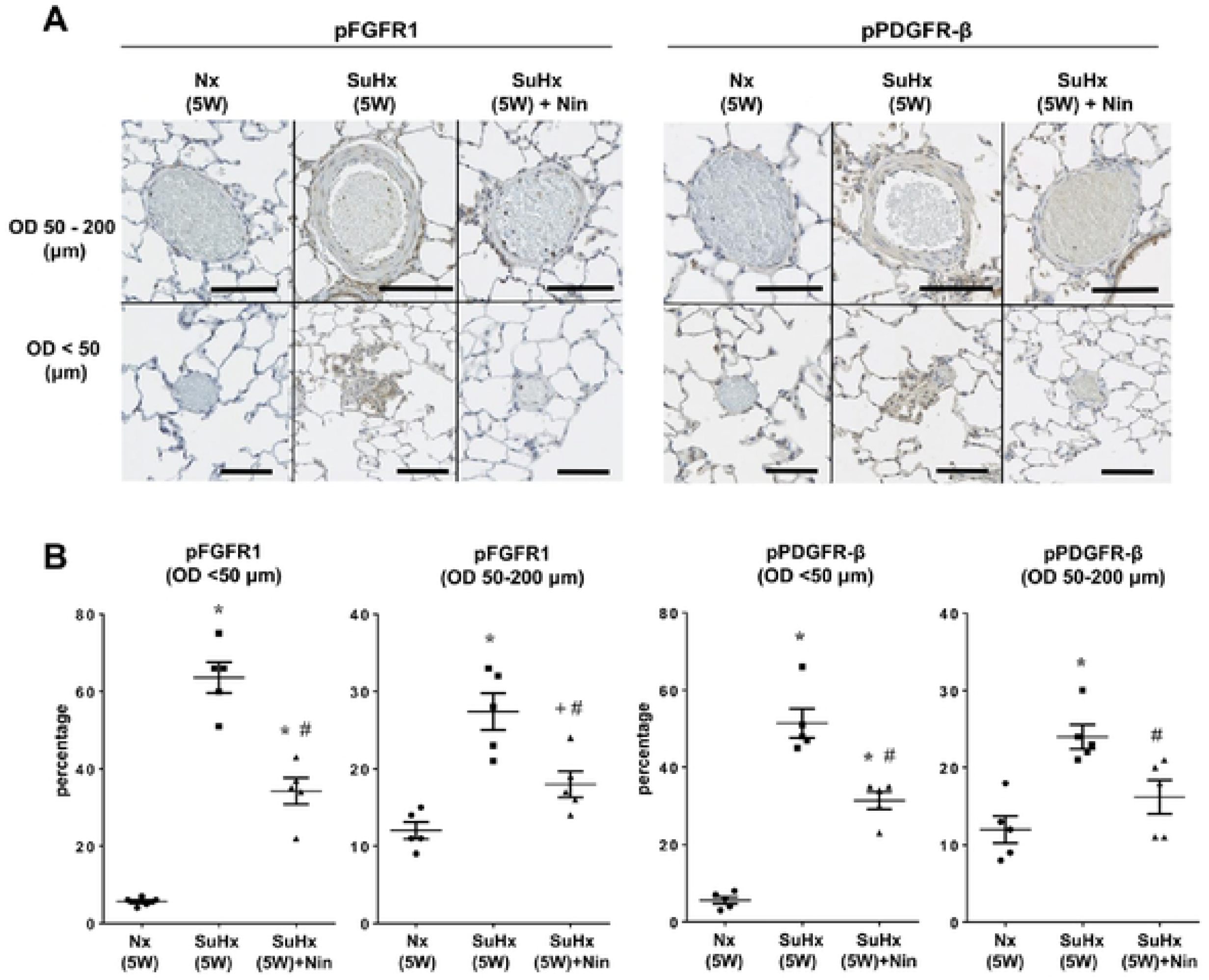
Expression of FGF and PDGF receptors in medial and neointimal lesions of small pulmonary arteries. (A) Representative immunohistochemistry of phosphorylated fibroblastic growth factor receptor 1 (pFGFR1) (left side) and phosphorylated platelet derived growth factor receptor-β (pPDGFR-β) (right side) in pulmonary arteries (200×). Scale bars indicate 50 μm (OD 50-200) and 10 μm (OD < 50). (B) Quantitative analysis of pFGFR1 and pPDGFR-β levels in pulmonary normotensive control rats at 5 weeks [Nx(5W)], Sugen 5416/hypoxic rats at 5 weeks [SuHx(5W)] after Sugen 5416 injection, and SuHx rats with nintedanib treatment from weeks 3 to 5 [SuHx(5W)+Nin]. Pulmonary arteries were divided into 50 – 200 μm outer diameter (OD) and < 50 μm OD for analysis. Plotted values are means ± SE (n=5). * p<0.0001 vs. Nx. + p<0.01 vs. Nx. # p<0.05 vs. SuHx(5W).

### Expression of Twist1 in rat lung tissue

The expression of Twist1 protein by western blotting was significantly augmented in lung tissue from SuHx(5W) rats compared with that from Nx(5W) rats, and nintedanib remarkably reduced this augmented expression (Figure 6).

**Figure 6.**
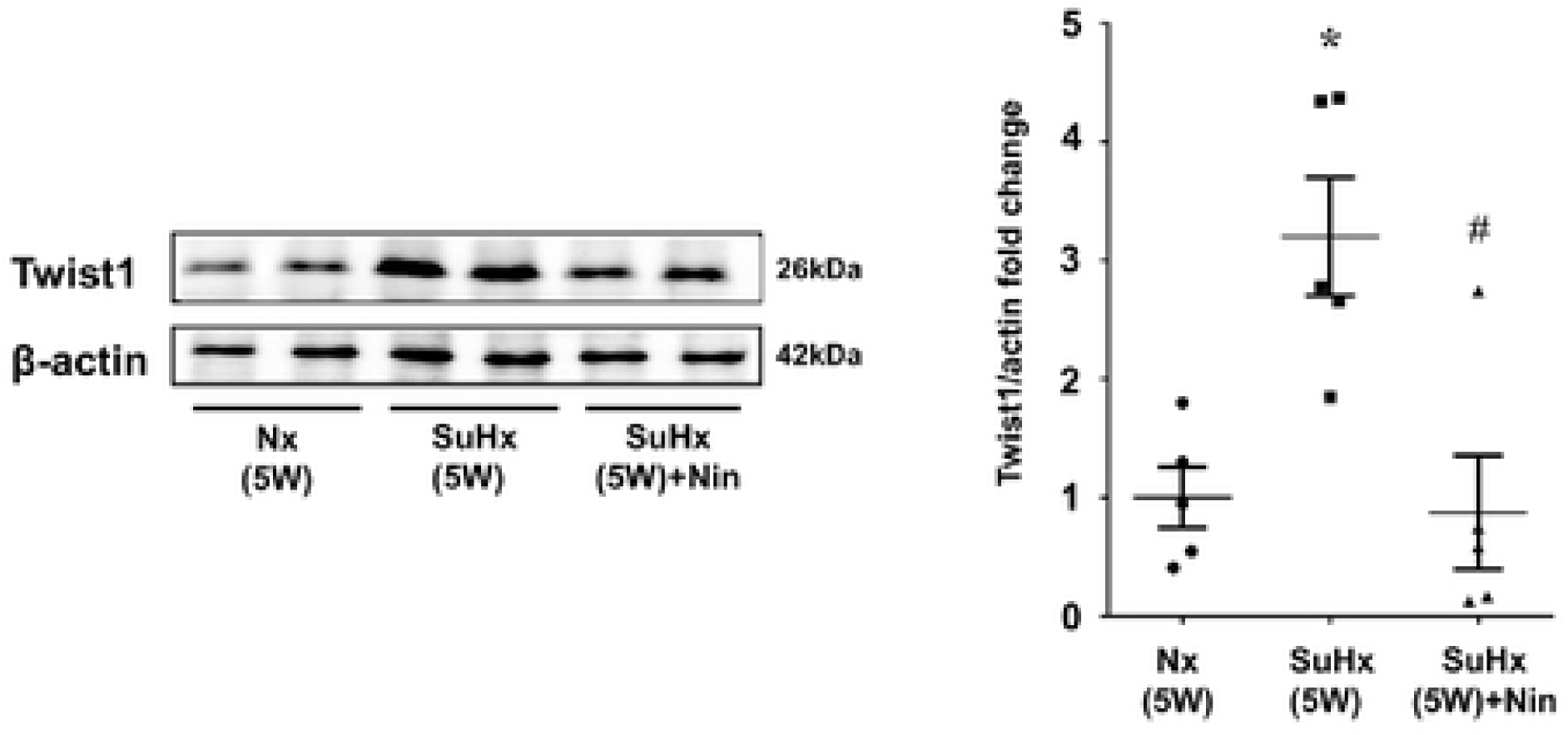
Expression of Twist1 protein in lung tissue of rats. Expression of Twist1 protein by western blotting analysis in lung tissues from pulmonary normotensive control rats at 5 weeks (Nx), Sugen 5416/hypoxic rats at 5 weeks [SuHx(5W)] after Sugen 5416 injection, and SuHx rats with nintedanib treatment from weeks 3 to 5 [SuHx(5W) + Nin]. Representative blots are shown (left). Protein signals were quantified by densitometric analysis, normalized to the corresponding β-actin signal, and plotted as fold changes (right). The plotted values shown are means ± SE (n=5). * p<0.001 vs. CON. # p<0.05 vs. SuHx(5W).

## Discussion

This report assessed the beneficial effect of nintedanib for PAH both *in vitro* and *in vivo*. Nintedanib attenuated the progression of EndMT in HPMVECs and inhibited the proliferation of HPASMCs *in vitro*. In addition, we demonstrated that phosphorylated PDGF and FGF receptors were increased in vascular occlusive neointimal lesions and the thickening medial wall in PAH rats. Two-week treatment of PAH rats with nintedanib significantly ameliorated pulmonary hemodynamics accompanied with improved vascular remodeling of PAs.

### Mechanism of pulmonary arterial remodeling in PAH

Medial wall thickening and occlusive neointimal lesion of PAs are common pathological findings in PAH. In the remodeling process of PAs in PAH, medial wall thickening driven by proliferation and hypertrophy of smooth muscle cells occurs early in disease progression. Moreover, the contribution of functional changes in smooth muscle cells of different phenotypes occurs particularly in the establish phase (24). On the other hand, a greater contribution of neointimal lesions than of medial wall thickness in the development of PAH was shown in an experimental model (14), and the role of EndMT in the progression of neointimal lesions has been recently suggested (10–13).

The essential role of EndMT in the pathogenesis of various cardiovascular diseases and PAH has been recently demonstrated (25). Occlusive neointimal lesions in small PAs, including plexiform lesions, are characteristic in PAH, and the proliferative cells that comprise these PAH-specific neointimal lesions have both endothelial and mesenchymal phenotypes (9, 11). Inducers, such as a hemodynamic stress, mechanical injury, hypoxia, and inflammation, upregulate several growth factors and cytokines, including PDGF, FGF, and TGF-β, which play important roles in the progression of EndMT (26, 27). Endothelial cells lose their intercellular adhesion after exposure to these stimuli, and subsequently change from a cobblestone to spindle-shaped morphology. EndMT-induced endothelial cells express both endothelial and mesenchymal markers, and acquire further capacity to proliferate, migrate, and avoid apoptosis (10). In a PAH model, both endothelial and mesenchymal markers were seen by immunohistochemistry in neointimal lesions of small PAs (12). Recent reports also have shown that green fluorescent protein (GFP)-labeled endothelial cells in mice were transformed by Sugen 5416 injection and chronic hypoxic exposure to a mesenchymal phenotype with high proliferative and migratory abilities (13). In human PAH, Ranchoux *et al*. demonstrated by electron microscopy that caveolae, which are characteristic of vascular endothelial cells, and dense bodies, which are dominant in vascular smooth muscle cells, were present in the same cells comprising occlusive neointimal lesions. Moreover, the expression of Twist1, a transcription factor associated with EndMT, was higher in human PAH lung than in normal human lung (11). These data indicate a critical contribution of EndMT in the progression of neointimal lesions in PAH.

### Tyrosine kinase inhibitors for PAH

Imatinib is a TKI for Bcr-Abl, c-Kit, c-Abl, and PDGF receptor signaling, and has been used for the treatment of chronic myelogenous leukemia and gastrointestinal stromal tumors (28). Imatinib was also expected to be a novel anti-vascular treatment for PAH due to its inhibitory effect on PDGF receptor. Imatinib was shown to inhibit the proliferation of HPASMCs from PAs in human PAH (29), and long-term treatment with imatinib prevented the development of chronic hypoxia- and monocrotaline-induced pulmonary hypertension in rats (30, 31). Furthermore, imatinib combined with various vasodilators significantly improved 6-minute walking distance and pulmonary vascular resistance in human patients, and a phase III clinical study of imatinib for PAH was implemented (32). However, imatinib did not receive final approval for PAH because of its serious adverse effects. Dasatinib, a TKI for Bcr-Abl, c-kit, Src family and PDGF receptor signaling, has been used also to treat chronic myelogenous leukemia. Importantly, dasatinib-induced pulmonary hypertension has been previously reported, suggesting specific toxicity of dasatinib on pulmonary vessels (33, 34). Nintedanib is a TKI for PDGF, FGF, and VEGF receptors. FGFR1, 2, and 3; PDGF receptor-α and -β; and VEGF receptor-1, −2, and −3 are receptors that are targeted by nintedanib (35). Nintedanib was originally developed for the treatment of malignancies as an agent to inhibit angiogenesis, cell proliferation, and migration (36). Nintedanib also reduced the mRNA and protein expression of extracellular matrix components in fibroblasts from human IPF (37), indicating the ability of nintedanib to treat IPF. Additionally, a recent phase III clinical study for human IPF showed beneficial effects of nintedanib in improving forced vital capacity and reducing the incidence of acute exacerbations. Diarrhea and liver dysfunction are major adverse effects of nintedanib, but these events were generally well tolerated (16). Hence, nintedanib has been approved globally for the treatment of IPF. In contrast to dasatinib, the results from the phase III study and postmarketing surveillance revealed no evidence to warrant elevating the risk of pulmonary hypertension by nintedanib. Based on this background, we hypothesized that nintedanib was a novel and tolerable treatment for PAH because of its anti-vascular remodeling via inhibition of PDGF and FGF signaling.

### Effect of nintedanib in vitro

In this study, nintedanib significantly prevented the increased expression of mesenchymal markers in EndMT-induced HPMVECs. A recent report showed that nintedanib reduced TGF-β signaling via inhibition of SMAD3 and p38 MAPK pathways (37). TGF-β signaling is considered a major regulator of EndMT, suggesting that the inhibition of TGF-β signaling is one of the mechanisms of the anti-EndMT effect by nintedanib. Moreover, previous reports showed that PDGF and FGF enhanced signaling of p38/AKT and PI3K/AKT pathways. Increased p38/PI3K/AKT signaling affected the expression of endothelial markers involving vWF and CD31 via transcription factors involving Twist1 (25). These data suggest that nintedanib can also regulate EndMT via inhibition of PDGF/FGF receptor tyrosine kinase activity in addition to TGF-β signaling. On the other hand, the reduced expression of endothelial markers was not restored after treatment with nintedanib. HPMVECs were pretreated with nintedanib before induction of EndMT in this study. Blockade of VEGF signaling by nintedanib may disrupt the maintenance of endothelial cell characteristics, because VEGF signaling generally plays an endothelial protective role (38).

In HPASMCs, nintedanib also significantly prevented the proliferation induced by multiple growth factors, and the inhibitory effect of nintedanib was significantly greater than that of imatinib after co-stimulation with multiple growth factors. The LDH assay showed no difference even after treatment with nintedanib, suggesting no toxicity of nintedanib for HPASMCs. ERK1/2 and AKT, which are downstream effectors of PDGF and FGF signaling, play roles in the regulation of vascular smooth muscle cell proliferation and apoptosis. Nintedanib also remarkably reduced the phosphorylation of ERK1/2 and AKT after co-stimulation, and this inhibitory effect of nintedanib was greater than that of imatinib. These *in vitro* results suggest a beneficial, anti-remodeling effect of nintedanib in the intima and media of PAs in PAH.

### Effect of nintedanib in vivo

Classical animal models of pulmonary hypertension that are induced by chronic hypoxic exposure and monocrotaline have no neointimal lesions in the small PAs. Hence, these models were considered inappropriate for the study of PAH. In the present study, we used SuHx-PAH rats, which have neointimal lesions similar to those in human PAH (39), to evaluate the effect of nintedanib. Toba *et al*. provided a detailed evaluation of the time course progression of vascular remodeling in the SuHx-PAH rat (14). They observed thickened medial walls and neointimal lesions in PAs from 3 to 5 weeks, whereas fibrotic vasculopathy involving plexiform lesions progressed 8 weeks following injection of Sugen 5416. Although the medial wall thickness decreased to a normal range during the late phase of PAH even with an elevated RVSP, the increased density of severely occlusive PAs was obviously correlated with RVSP elevation. Thus, the authors concluded that the contribution of medial wall thickness was transient in the early phase of PAH, and the role of neointimal lesions in the small PAs seemed to be greater in the progression of disease. Whereas neointimal lesions are an important therapeutic target for the treatment of PAH, vascular lesions with substantial fibrosis are generally considered refractory to treatment (40). Therefore, we tested the anti-remodeling effect of nintedanib from 3 to 5 weeks in this study, targeting non-plexiform cellular neointimal vascular lesions. The phosphorylation of PDGF and FGF receptors significantly increased in the proliferative neointimal lesions and medial walls in the PAH rat, indicating that these PAH-specific neointimal lesions could be a major therapeutic target of nintedanib. In fact, nintedanib treatment from 3 to 5 weeks significantly reduced the expression of those receptors and prevented the progression of neointimal lesions, resulting in improved pulmonary hemodynamics in the PAH rats. Furthermore, we showed increased expression of Twist1 protein in the lung tissue from SuHx-PAH rats. Twist1 enhances the expression of TGF-β receptor and phosphorylation of SMAD2, and can lead to EndMT in PA endothelial cells (41). Chronic nintedanib significantly reduced Twist1 protein expression in this study. These findings indicate not only the contribution of EndMT in the development of proliferative occlusive neointimal lesions in PAH rats, but also the anti-EndMT effects of nintedanib. Additionally, it has been reported that BIBF1000, a structural precursor of nintedanib, did not disturb RV function in rats subjected to mechanical pressure overload by pulmonary artery-banding (42). Another recent study also showed that less dilatation, decreased fibrosis and hypertrophy of right ventricle after chronic treatment with nintedanib in Su/Hx-PAH rat by echocardiography and histological analysis (43), suggesting no cardiac toxicity of nintedanib.

A recently published study demonstrated that nintedanib had no therapeutic effect on rat and human PAH (44), which contradicted our results. Several factors could explain the discrepancy between the studies. For instance, rat strain that were used for experimental PAH were different. A previous report has shown that the severity of the response to Sugen5416 with hypoxic exposure was remarkably different between rat strains (45). Actually, the RVSP of SuHx PAH in Wistar-Kyoto rats was obviously lower in a recent study than in this study. Therefore, the response to nintedanib also may be dissimilar between strains. As another factor, muscularization of the small PAs was evaluated to assess the effect of nintedanib in a recent study. Such measurement is common for hypoxic or monocrotaline-induced pulmonary hypertension models that have no neointimal lesions, but it is not appropriate for the SuHx PAH rat which has neointimal lesions in the small PAs. In addition, pathological images of small PAs were not shown in that study; thus, it is unclear which parts of the small PAs were evaluated. Furthermore, the delivery route of nintedanib to the rat was not mentioned. Thus, the possibility remain that differences in the administration route of nintedanib may be the cause of the contradictory results. In contrast, Huang et al. reported that chronic treatment with nintedanib reduced pulmonary vascular remodeling in the Fos-related antigen-2 mouse model of systemic sclerosis (46), which was supportive of our results. On the other hand, another more recent study showed that chronic treatment with nintedanib of established late-phase PAH, from 8 to 11 weeks, did not improved pulmonary hemodynamics and vascular remodeling in SuHx rats (43). The results of these recent studies of established animal and human PAH suggest that the therapeutic power of nintedanib may be limited at least in advanced stages of PAH. Although the efficacy of nintedanib for mild to moderate PAH is still uncertain, the therapeutic benefit of nintedanib may rest with its ability to augment the effect of other vasodilators or to prevent the progression of PAH.

## Limitations

Our study has a few limitations. First, we have not yet assessed the phenotype of HPASMCs after stimulation; investigations into the effect of nintedanib on the functional changes in HPASMCs may bring further beneficial information. Second, while HPASMC used in this study is mainly isolated from the proximal PA, contribution of small distal PA is greater than that of large proximal PA in the disease progression of PAH. Thus, data would be more useful by using HPASMC isolated from only small distal PA. Third, the mechanism of the inhibitory effect of nintedanib on EndMT progression is still unclear. We induced EndMT in HPMVECs by using three different inducers simultaneously; nintedanib could have inhibited all three signaling pathways. The inhibitory mechanism was probably very complex and beyond the scope of the current study. Fourth, higher concentrations or longer regimens of nintedanib treatment for PAH in the rat have not yet been evaluated. Further experiments are necessary in the future to evaluate these limitations of our study.

## Conclusion

In conclusion, we have shown the beneficial effects of nintedanib for PAH *in vitro* and *in vivo*. Chronic treatment with nintedanib reversed the elevated pulmonary arterial pressure in the PAH rat via anti-EndMT effects in HPMVECs and anti-proliferative effects in HPASMCs. Moreover, an expanded indication of nintedanib for the treatment of PAH would be advantageous, since nintedanib has been approved already for the treatment of other human diseases with good tolerability. Although further investigations are necessary, the results of this study indicate that nintedanib may be a novel additional option for the treatment of human PAH, with an anti-vascular remodeling effect.

## Acknowledgments

We thank Hiroshi Kawai and Reiko Mineki for their technical assistance.

## Author contributions

T.T contributed whole of experiment. T.N supervised whole of this study. T.Y, L.W and S.K assisted in molecular biological experiment. Y.S and Y.N assisted in animal experiment. N.H, Y.K, F.T, Y.M, and K.T assisted in planning of this study.

## Conflict of interest

none declared.

## Funding

This work was supported by Grants-in-Aid for Scientific Research (C) from the Japan Society for the Promotion of Science (25461197).

## References

1. Lai YC, Potoka KC, Champion HC, Mora AL, Gladwin MT. Pulmonary arterial hypertension: the clinical syndrome. Circ Res. 2014;115(1):115–30.

2. Hurdman J, Condliffe R, Elliot CA, Davies C, Hill C, Wild JM, et al. ASPIRE registry: assessing the Spectrum of Pulmonary hypertension Identified at a REferral centre. Eur Respir J. 2012;39(4):945–55.

3. Frost AE, Badesch DB, Miller DP, Benza RL, Meltzer LA, McGoon MD. Evaluation of the predictive value of a clinical worsening definition using 2-year outcomes in patients with pulmonary arterial hypertension: a REVEAL Registry analysis. Chest. 2013;144(5):1521–9.

4. Noskovicova N, Petrek M, Eickelberg O, Heinzelmann K. Platelet-derived growth factor signaling in the lung. From lung development and disease to clinical studies. Am J Respir Cell Mol Biol. 2015;52(3):263–84.

5. Perros F, Montani D, Dorfmuller P, Durand-Gasselin I, Tcherakian C, Le Pavec J, et al. Platelet-derived growth factor expression and function in idiopathic pulmonary arterial hypertension. Am J Respir Crit Care Med. 2008;178(1):81–8.

6. Benisty JI, McLaughlin VV, Landzberg MJ, Rich JD, Newburger JW, Rich S, et al. Elevated basic fibroblast growth factor levels in patients with pulmonary arterial hypertension. Chest. 2004;126(4):1255–61.

7. Tu L, Dewachter L, Gore B, Fadel E, Dartevelle P, Simonneau G, et al. Autocrine fibroblast growth factor-2 signaling contributes to altered endothelial phenotype in pulmonary hypertension. Am J Respir Cell Mol Biol. 2011;45(2):311–22.

8. Izikki M, Guignabert C, Fadel E, Humbert M, Tu L, Zadigue P, et al. Endothelial-derived FGF2 contributes to the progression of pulmonary hypertension in humans and rodents. J Clin Invest. 2009;119(3):512–23.

9. Tuder RM, Groves B, Badesch DB, Voelkel NF. Exuberant endothelial cell growth and elements of inflammation are present in plexiform lesions of pulmonary hypertension. The American journal of pathology. 1994;144(2):275–85.

10. Arciniegas E, Frid MG, Douglas IS, Stenmark KR. Perspectives on endothelial-to-mesenchymal transition: potential contribution to vascular remodeling in chronic pulmonary hypertension. Am J Physiol Lung Cell Mol Physiol. 2007;293(1):L1–8.

11. Ranchoux B, Antigny F, Rucker-Martin C, Hautefort A, Pechoux C, Bogaard HJ, et al. Endothelial-to-mesenchymal transition in pulmonary hypertension. Circulation. 2015;131(11):1006–18.

12. Good RB, Gilbane AJ, Trinder SL, Denton CP, Coghlan G, Abraham DJ, et al. Endothelial to Mesenchymal Transition Contributes to Endothelial Dysfunction in Pulmonary Arterial Hypertension. The American journal of pathology. 2015;185(7):1850–8.

13. Suzuki T, Carrier EJ, Talati MH, Rathinasabapathy A, Chen X, Nishimura R, et al. Isolation and characterization of endothelial-to-mesenchymal transition cells in pulmonary arterial hypertension. Am J Physiol Lung Cell Mol Physiol. 2018;314(1):L118–L26.

14. Toba M, Alzoubi A, O’Neill KD, Gairhe S, Matsumoto Y, Oshima K, et al. Temporal hemodynamic and histological progression in Sugen5416/hypoxia/normoxia-exposed pulmonary arterial hypertensive rats. Am J Physiol Heart Circ Physiol. 2014;306(2):H243–50.

15. Wollin L, Wex E, Pautsch A, Schnapp G, Hostettler KE, Stowasser S, et al. Mode of action of nintedanib in the treatment of idiopathic pulmonary fibrosis. Eur Respir J. 2015;45(5):1434–45.

16. Richeldi L, du Bois RM, Raghu G, Azuma A, Brown KK, Costabel U, et al. Efficacy and safety of nintedanib in idiopathic pulmonary fibrosis. N Engl J Med. 2014;370(22):2071–82.

17. Nagai T, Kanasaki M, Srivastava SP, Nakamura Y, Ishigaki Y, Kitada M, et al. N-acetyl-seryl-aspartyl-lysyl-proline inhibits diabetes-associated kidney fibrosis and endothelial-mesenchymal transition. Biomed Res Int. 2014;2014:696475.

18. Zhou H, Wang Y, Zhou Q, Wu B, Wang A, Jiang W, et al. Down-Regulation of Protein Kinase C-epsilon by Prolonged Incubation with PMA Inhibits the Proliferation of Vascular Smooth Muscle Cells. Cell Physiol Biochem. 2016;40(1-2):379–90.

19. Muir D, Varon S, Manthorpe M. An enzyme-linked immunosorbent assay for bromodeoxyuridine incorporation using fixed microcultures. Anal Biochem. 1990;185(2):377–82.

20. Wakatsuki S, Furuno A, Ohshima M, Araki T. Oxidative stress-dependent phosphorylation activates ZNRF1 to induce neuronal/axonal degeneration. J Cell Biol. 2015;211(4):881–96.

21. Sakao S, Tatsumi K. The effects of antiangiogenic compound SU5416 in a rat model of pulmonary arterial hypertension. Respiration. 2011;81(3):253–61.

22. Kuriyama S, Morio Y, Toba M, Nagaoka T, Takahashi F, Iwakami S, et al. Genistein attenuates hypoxic pulmonary hypertension via enhanced nitric oxide signaling and the erythropoietin system. Am J Physiol Lung Cell Mol Physiol. 2014;306(11):L996–L1005.

23. Pullamsetti SS, Berghausen EM, Dabral S, Tretyn A, Butrous E, Savai R, et al. Role of Src tyrosine kinases in experimental pulmonary hypertension. Arteriosclerosis, thrombosis, and vascular biology. 2012;32(6):1354–65.

24. Stenmark KR, Frid MG, Graham BB, Tuder RM. Dynamic and diverse changes in the functional properties of vascular smooth muscle cells in pulmonary hypertension. Cardiovasc Res. 2018;114(4):551–64.

25. Jackson AO, Zhang J, Jiang Z, Yin K. Endothelial-to-mesenchymal transition: A novel therapeutic target for cardiovascular diseases. Trends Cardiovasc Med. 2017;27(6):383–93.

26. Li Z, Jimenez SA. Protein kinase Cdelta and c-Abl kinase are required for transforming growth factor beta induction of endothelial-mesenchymal transition in vitro. Arthritis Rheum. 2011;63(8):2473–83.

27. Song S, Zhang M, Yi Z, Zhang H, Shen T, Yu X, et al. The role of PDGF-B/TGF-beta1/neprilysin network in regulating endothelial-to-mesenchymal transition in pulmonary artery remodeling. Cell Signal. 2016;28(10):1489–501.

28. Peng B, Lloyd P, Schran H. Clinical pharmacokinetics of imatinib. Clinical pharmacokinetics. 2005;44(9):879–94.

29. Nakamura K, Akagi S, Ogawa A, Kusano KF, Matsubara H, Miura D, et al. Pro-apoptotic effects of imatinib on PDGF-stimulated pulmonary artery smooth muscle cells from patients with idiopathic pulmonary arterial hypertension. Int J Cardiol. 2012;159(2):100–6.

30. Ciuclan L, Hussey MJ, Burton V, Good R, Duggan N, Beach S, et al. Imatinib attenuates hypoxia-induced pulmonary arterial hypertension pathology via reduction in 5-hydroxytryptamine through inhibition of tryptophan hydroxylase 1 expression. Am J Respir Crit Care Med. 2013;187(1):78–89.

31. Pankey EA, Thammasiboon S, Lasker GF, Baber S, Lasky JA, Kadowitz PJ. Imatinib attenuates monocrotaline pulmonary hypertension and has potent vasodilator activity in pulmonary and systemic vascular beds in the rat. Am J Physiol Heart Circ Physiol. 2013;305(9):H1288–96.

32. Hoeper MM, Barst RJ, Bourge RC, Feldman J, Frost AE, Galie N, et al. Imatinib mesylate as add-on therapy for pulmonary arterial hypertension: results of the randomized IMPRES study. Circulation. 2013;127(10):1128–38.

33. Montani D, Bergot E, Gunther S, Savale L, Bergeron A, Bourdin A, et al. Pulmonary arterial hypertension in patients treated by dasatinib. Circulation. 2012;125(17):2128–37.

34. Guignabert C, Phan C, Seferian A, Huertas A, Tu L, Thuillet R, et al. Dasatinib induces lung vascular toxicity and predisposes to pulmonary hypertension. J Clin Invest. 2016;126(9):3207–18.

35. Inomata M, Nishioka Y, Azuma A. Nintedanib: evidence for its therapeutic potential in idiopathic pulmonary fibrosis. Core Evid. 2015;10:89–98.

36. Caglevic C, Grassi M, Raez L, Listi A, Giallombardo M, Bustamante E, et al. Nintedanib in non-small cell lung cancer: from preclinical to approval. Ther Adv Respir Dis. 2015;9(4):164–72.

37. Rangarajan S, Kurundkar A, Kurundkar D, Bernard K, Sanders YY, Ding Q, et al. Novel Mechanisms for the Antifibrotic Action of Nintedanib. Am J Respir Cell Mol Biol. 2016;54(1):51–9.

38. Voelkel NF, Gomez-Arroyo J. The role of vascular endothelial growth factor in pulmonary arterial hypertension. The angiogenesis paradox. Am J Respir Cell Mol Biol. 2014;51(4):474–84.

39. Abe K, Toba M, Alzoubi A, Ito M, Fagan KA, Cool CD, et al. Formation of plexiform lesions in experimental severe pulmonary arterial hypertension. Circulation. 2010;121(25):2747–54.

40. Sakao S, Tatsumi K, Voelkel NF. Reversible or irreversible remodeling in pulmonary arterial hypertension. Am J Respir Cell Mol Biol. 2010;43(6):629–34.

41. Kwapiszewska G, Crnkovic S, Stenmark KR. A Twist on Pulmonary Vascular Remodeling: Endothelial to Mesenchymal Transition? Am J Respir Cell Mol Biol. 2018;58(2):140–1.

42. de Raaf MA, Herrmann FE, Schalij I, de Man FS, Vonk-Noordegraaf A, Guignabert C, et al. Tyrosine kinase inhibitor BIBF1000 does not hamper right ventricular pressure adaptation in rats. Am J Physiol Heart Circ Physiol. 2016;311(3):H604–12.

43. Rol N, de Raaf MA, Sun X, Kuiper VP, da Silva Goncalves Bos D, Happe C, et al. Nintedanib improves cardiac fibrosis but leaves pulmonary vascular remodeling unaltered in experimental pulmonary hypertension. Cardiovasc Res. 2018.

44. Richter MJ, Ewert J, Grimminger F, Ghofrani HA, Kojonazarov B, Petrovic A, et al. Nintedanib in Severe Pulmonary Arterial Hypertension. Am J Respir Crit Care Med. 2018.

45. Jiang B, Deng Y, Suen C, Taha M, Chaudhary KR, Courtman DW, et al. Marked Strain-Specific Differences in the SU5416 Rat Model of Severe Pulmonary Arterial Hypertension. Am J Respir Cell Mol Biol. 2016;54(4):461–8.

46. Huang J, Maier C, Zhang Y, Soare A, Dees C, Beyer C, et al. Nintedanib inhibits macrophage activation and ameliorates vascular and fibrotic manifestations in the Fra2 mouse model of systemic sclerosis. Ann Rheum Dis. 2017;76(11):1941–8.

